# Poor sensing maximises microbial fitness when few out of many signals are sensed

**DOI:** 10.1101/800292

**Authors:** Age J. Tjalma, Robert Planqué, Frank J. Bruggeman

**Affiliations:** Systems Biology Lab, AIMMS, VU University, De Boelelaan 1085, 1081 HV, Amsterdam, The Netherlands; Dept. of Mathematics, VU University, De Boelelaan 1111, 1081 HV, Amsterdam, The Netherlands

**Keywords:** Cellular decision making, Phenotypic adaptation strategies, Bethedging, Cellular signalling, Information Theory, Mutual information, Biochemical noise

## Abstract

An open problem in biology is to understand when particular adaptation strategies of microorganisms are selected during evolution. They range from random, bet-hedging strategies to deterministic, responsive strategies, relying on signalling circuits. We present an evolutionary model that integrates basic statistical physics of molecular circuits with fitness maximisation and information theory. This model provides an explanation for a puzzling observation on responsive strategies: the accuracy with which signalling networks track external signals seems remarkably low. Single cells often distinguish only between 2 to 4 concentration ranges, corresponding to 1 or 2 bits of mutual information between signal and response. Why did evolution lead to such low-fidelity signalling systems? Our theory offers an explanation by taking a novel perspective. It considers the fitness benefit of all signals, including those that are not sensed. We introduce a new concept, ‘latent information’, which captures the mutual information between all non-sensed signals and the optimal response. The theory predicts that it is often evolutionarily optimal to transduce sensed signals noisily when latent information is present. It indicates that fitness can indeed be maximal when the mutual information extracted from sensed signals is not maximal, but rather has a low value of about 1 or 2 bits. Cells likely do not sense all signals because of the fitness cost of expressing idle signalling systems that consume limited biosynthetic resources. Our theory illustrates that as the total available information about the optimal behaviour decreases, the cell should trust the available information less, and gamble more.

**Significance Statement:** Surprisingly, microorganisms appear to sense only very few environmental signals (such as nutrients and stresses) compared to the number of conditions they can encounter. Even worse, the signals they do sense are transduced at low fidelity. We study the accuracy of sensing in the situation where multiple signals determine the optimal response, but only few signals are actually sensed. We show that it is in fact to be expected that sensing in the presence of latent information should be imprecise, and that signalling circuits should underperform.

---

Evolutionary theories about the fitness of phenotypic strategies (1–6) have made successful predictions of the outcomes of natural selection in fluctuating environments (3, 4, 7, 8). In those theories, the (geometric) fitness (4–6) is generally maximised. This fitness measure equals the logarithm of the fold change in the number of organisms divided by the number of environmental periods, or the total time. Organisms that produced most offspring have maximal geometric fitness. These theories evaluate the fitness contributions of phenotypic strategies, ranging from random, bet-hedging strategies to deterministic, responsive strategies.

Bet-hedging strategies (14) are independent of environmental conditions. Cells can switch randomly between alternative phenotypic states. This ensures the existence of subpopulations that might be maladapted to the current condition, but are prepared for future conditions. For example, slow-growing, stress-tolerant cells (so-called persister cells) and fast-growing, stress-sensitive cells have been discovered in microbial cell populations (9–12). The fast growing cells largely determine the current fitness, while the persister cells are ‘insurance policies’, guaranteeing survival when conditions suddenly become harsh and extinction-threatening. Together, they maximise the geometric fitness of the population in conditions that fluctuate between benign and adverse conditions.

Close-to-deterministic adaptation strategies, on the other hand, also exist. Cells then perceive and transmit a signal to infer the state of the environment and adapt their phenotype accordingly. The omnipresence of two-component signalling circuits across microorganisms (13) highlights the importance of this mode of adaptation. However, inevitable molecular stochasticity in signalling circuits can cause random mismatches between the phenotype and the environmental state, leading to fitness losses.

Since the entire range of phenotypic behaviours is found amongst microorganisms, and because they are great experimental systems for physiology and evolution, microbiology is the perfect playground for testing predictions and improving evolutionary theories. We therefore focus our theory on microbial phenotypic adaptation.

The question under which conditions bet-hedging adaptation strategies are more evolutionarily successful than sensing strategies has a long history in evolutionary biology. Bet-hedging has been predicted to be evolutionarily advantageous under at least two conditions (14): In slowly-changing, mild environments where sensing machinery would rarely provide an evolutionary benefit for long periods of time, and in environments that change quickly into extinction-threatening states where responsive adaptation would be too slow (1–4, 6, 14). In all other cases, responsive signalling strategies should be favoured.

What remains poorly understood is what the optimal accuracy of signalling systems should be. In particular, experiments indicate that single cells have a remarkably low capacity for accurate tracking of environmental signals (15–19) far below theoretical predictions: cells should be able to measure much more accurately than they actually do. Intuitively, one would expect that more accurate signalling improves evolutionary success as it reduces maladaptation. Accordingly, many theories are based on maximisation of the mutual information between signal and response (20–22). However, in reality cells only appear to sense few external signals and in a noisy regime, leading to stochastic responses (23–26). Moreover, experiments report poor signalling capabilities of single cells; no more than 2 bits of mutual information between an external signal and an internal response have been found (16–19). Thus, either reliable signal transduction is not that important for evolutionary success, or it is very hard for cells to achieve reliable signal transduction.

Reliable signal transduction might be difficult to achieve due to the inevitable randomness of molecular systems (27). Increased expression of signalling proteins can make them more reliable (28), but requires a fitness-reducing diversion of resources from growth processes. Only if signalling activity leads to a net fitness increase one would expect signalling systems to evolve. Therefore, expression of idle signalling systems is likely fitness reducing in the absence of the signal and, moreover, genes encoding signalling machinery may then eventually accumulate random mutations that reduce signalling performance. In addition, many signalling systems rely on signal receptors in the membrane, and their expression is at the expense of nutrient importers; this reduces fitness too. These considerations may explain why cells express so few signalling systems; *E. coli* can grow, for instance, on hundreds of carbon sources, but it only has a handful one- and two-component sensing systems dedicated to nutrient sensing. However, they do not explain why exploited signalling systems display such remarkably low mutual information values between their signal and their response. Answering this question is the goal of this paper.

In information-theoretic studies on cellular signalling, maximisation of mutual information between a signal and a response variable is considered beneficial for cells. We argue that this is not necessarily so. Imagine, for example, a flat relationship between the fitness value and the response variable versus one that is sharply peaked. In the former case, infinite mutual information between signal and response variable would not improve fitness, while it would in the latter case. More importantly, another limitation is that most studies consider only one signal and one response variable. It is, however, likely that cells integrate different signals to induce a single response. Imagine a fitness landscape as a function of several response variables that contains ridges and peaks. In such a landscape, mutual information maximisation with respect to one signal and one response variable is insufficient. Instead, phenotypic adaptation strategies should be evaluated on the basis of fitness maximisation—considering all signals. Importantly, also ‘latent signals’ – those that are not sensed by the cell, but that are fitness enhancing if they would be sensed – should be considered.

## Results

### The geometric fitness model

We aim to investigate fitness maximisation of an isogenic population in the presence of fluctuating signals. All single cells sense a signal *s* that induces the expression of a response protein with concentration *p* (figure 1A). In addition to the sensed signal, ‘latent’ signals, *s*_*l*1_, …, *s*_*ln*_, exist that are not sensed by cells. They are the elements of the vector *s*_*l*_. All signals, both sensed and latent, determine the optimal protein concentration *p*_*o*_(*s*, *s*_*l*_) that maximises fitness.

**Fig. 1.**
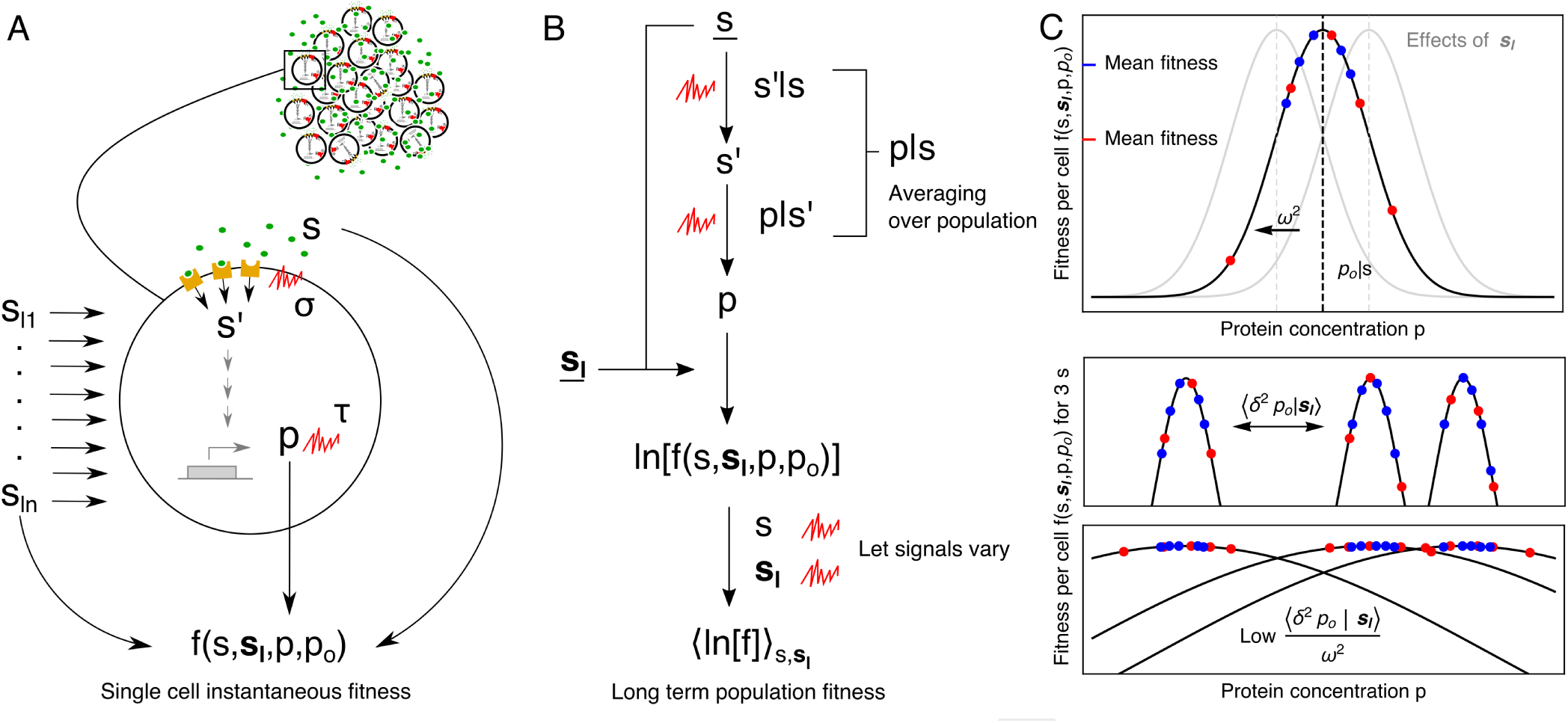
Biological situation, methodological workflow, and intuition for the fitness function. **A.** Evolution acts on an (isogenic) population. An individual cell senses an environmental signal *s* and estimates its concentration to be *s′*. In this process noise is added, with variance *σ*^2^. The signal is transduced and eventually leads to protein concentration *p*, which contains noise on its own; with variance *τ*^2^. The concentration *p* leads to a certain fitness of the cell depending on how far it is from the optimum *p*_*o*_ following the fitness function *f* (*s*, **s**_*l*_, *p*, *p*_*o*_). The optimal protein concentration *p*_*o*_ is a function of the sensed signal *s* and all latent signals *s*_*l*1_, …, *s*_*ln*_. **B.** Under a fixed environment we use 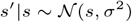 and 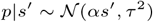 to obtain 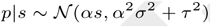, as shown in the appendix. The average fitness of the phenotypically heterogeneous population at one environment can now be determined and depends on *s*. Taking the logarithm and averaging over all signals (i.e., all environments) leads to the geometric mean of the fitness. **C.** The fitness of one cell is given by: 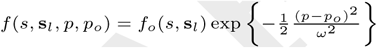 where *ω* determines the width of the fitness function, and *f*_*o*_(*s*, **s**_*l*_) determines the fitness (growth rate) in the optimum. The optimal protein concentration *p*_*o*_ is determined by the signal *s* (black curve), but is also influenced by the latent signals **s**_*l*_ (gray curves). Considering two hypothetical populations of blue and red cells, where the blue cells are genetically superior in transducing signals, we see how the mean fitness of the population in one environment depends on the accuracy of estimating the optimum *p*_*o*_. However, depending on the latent signals **s**_*l*_, the optimum and thus the mean fitness can still change. The middle plot shows how, under fixed **s**_*l*_, the variance in optima determines how far the peaks of the fitness curves lie apart. The bottom plot illustrates how inaccurate signal transduction will have fewer fitness consequences when the variance in optima is small relative to the width of the fitness curve.

We use the following fitness function, which reaches its maximal value *f*_*o*_(*s*, *s*_*l*_) at *p* = *p*_*o*_,

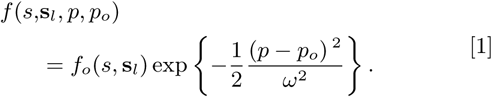

The fitness reduction, due to a deviation of the protein concentration *p* from its optimal value *p*_*o*_ is determined by the width of the “fitness-landscape” *ω* (figures 1A and 1C). The interpretation of the fitness equation is explained in figure 1C.

To keep the model analytically tractable, we assume in addition that: (1) the optimum *p*_*o*_ is given by the deterministic function *q*(*s*, *s*_*l*_), which is a nonlinear function of *s* plus a weighted sum of all latent signals s_*l*_ such that

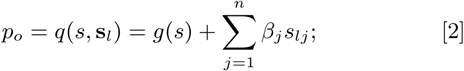

(2) all latent signals in s_*l*_ are independent, i.e., they do not covary, and are normally distributed.

Given these assumptions, following the methodology explained in figure 1B, we obtain for the fitness of a population of isogenic cells sensing one signal, averaged over all latent signals (see the appendix for the general derivation with multiple sensed signals),

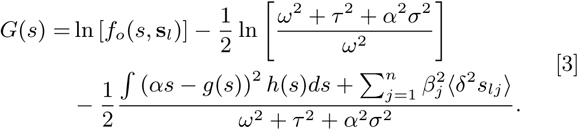

The first term ln[*f*_o_ (*s*, *s*_*l*_)] equals the maximal geometric fitness, we will denote it as ln[*fo*] from here on. The last two terms are fitness costs. The first captures the cost of noisy signalling and quantifies the fitness reduction due to protein fluctuations that lead to an average deviation from the optimal concentration. The total variance of the protein fluctuations equals *τ*^2^ + *α*^2^*σ*^2^. The second cost term captures a distance from optimality due to two suboptimal (deterministic) effects: i. the relationship 〈*p|s*〉 = *αs* might deviate from the optimal relation *g*(*s*) and ii. not sensing of latent signals that are informative about the value of *q*(*s*, s_*l*_), leads to a fitness loss too, in terms of the variances of these latent signals.

The last cost term in equation 3 contains *∫*(*αs* − *g*(*s*))^2^*h*(*s*)*ds*, which is reminiscent of a mean squared error (MSE) of an estimator. Its occurrence suggests that evolution can be interpreted as minimising the distance of a cell’s behaviour to its optimal behaviour. It is debatable whether this term can be made small by natural selection. For this to occur, the dependency *g*(*s*) should not become a too complex function, as otherwise cells would not be able to approximate it by a steady-state input-output relation of a molecular circuit.

Considering that we aim to investigate optimal sensing cells, we assume that *g*(*s*) ≈ *αs*. Therefore, the equation for the geometric mean fitness of sensing cells in the presence of multiple signals, both latent and observed, reduces to

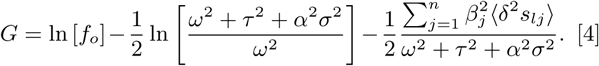

The final term in equation 4 is the fitness cost of varying latent signals. It quantifies how much fitness-influencing uncertainty remains about the environment, caused by the influence of all latent signals s_*l*_ on the optimum *p*_*o*_. This fitness cost is dependent on the width of the fitness function and the internal variance, and can thus be reduced by increased protein fluctuations, i.e., by increasing *τ*^2^ + *α*^2^*σ*^2^. This would lead to bet-hedging or ‘noisy sensing’ strategies, in agreement with previous results (1–3). Such strategies can be optimal, which is shown by differentiating equation 4 with respect to the internal variance and solving for its optimum value. Division by *ω*^2^ then gives,

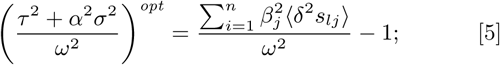

indicating that the relative internal variance should only exceed zero when the relative uncertainty in the environment exceeds 1 (figure 2A). In this case noisy signalling pays off due uncertainty in the environment. The geometric mean fitness of a cell is more sensitive to the relative uncertainty about its environment than to its own relative internal variance (figure 2A).

Thus, an optimal sensing cell, which does not sense all signals that are informative about the optimal behaviour, will still have to be noisy to overcome fitness variation due to the influences of all latent, non-sensed signals. Such a cell would therefore not follow a deterministic, pure sensing strategy, but allow for some bet-hedging behaviour, indicating that the mutual information between *s* and *p* should not always be maximised. This is an important insight from our theory that we will explore further.

**Fig. 2.**
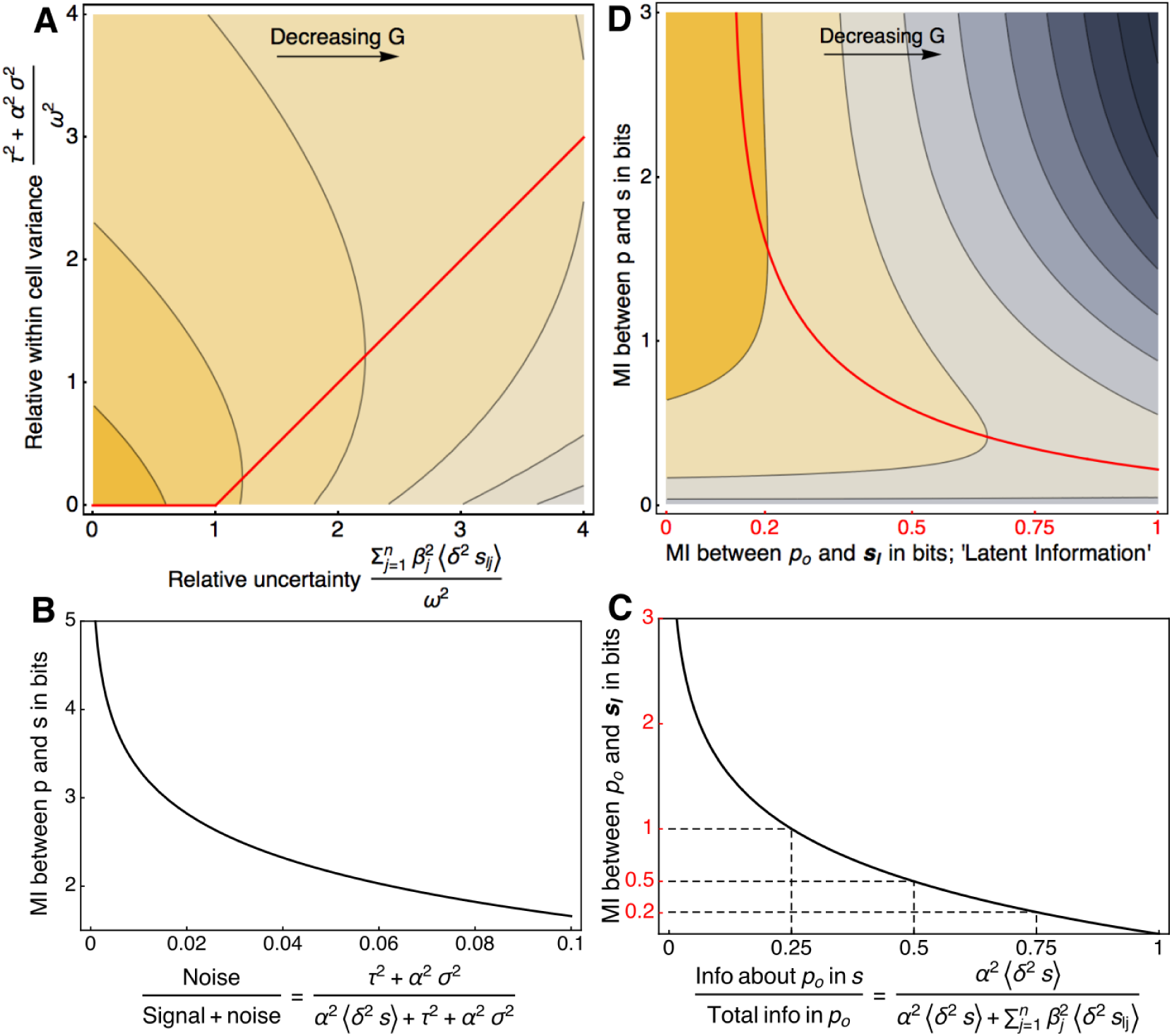
Mutual information, latent information and fitness of sensing cells. **A.** Contour plot of the geometric mean fitness (*G*) as a function of the relative within-cell variance and relative uncertainty in the environment. The colour indicates *G* as a percentage of the maximum (given by ln[*f*_*o*_]), here it ranges from 90% (bottom left) to 30% (bottom right). The red line shows the optimal relative internal variance for a given relative uncertainty. **B.** Mutual information (MI) between *p* and *s* as a function of the fraction of internal variance (i.e., noise) over all variance in *p*. This gives an indication of the range of the MI between *p* and *s* given a certain noise level. Note that the x-axis ranges from 0 to 0.1. **C.** MI between *p*_*o*_ and **s**_*l*_ as a function of the propagated variance in *p*_*o*_ from *s* over the total variance in *p*_*o*_. Since we are considering normal distributions, variance is proportional to mutual information. **D.** Contour plot of *G* as a function of the MI between the sensed signal *s* and the cell’s response *p* and the MI between the optimum *p*_*o*_ and all latent signals **s**_*l*_ (i.e., the latent information). The colouring indicates *G* as a percentage of the maximum; ranging from 75% (top left) to −150% (top right), the red line is the optimal MI between *p* and *s*, and *ρ* = 5. The x-axis covers a much smaller range than the y-axis, it corresponds to the red y-axis in C.

### Sensing vs non-sensing cells

The geometric mean fitness of a non-sensing population can also be derived from our general framework, outlined in the appendix. The variance induced by sensing machinery *σ*^2^ drops out of the equation, as non-sensing cells lack this machinery. However, since all signals are now latent, an extra variance term appears in the second cost term:

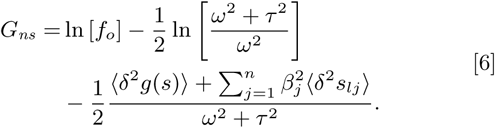

This extra variance term, 〈*δ*^2^*g*(*s*)〉, is the variance in the optimum due to the signal *s*, which is the signal of interest in the sensing population. Since we assumed above *g*(*s*) ≈ *αs*, we substitute 〈*δ*^2^*g*(*s*)〉 with *α*^2^〈*δ*^2^*s*〉. The optimal, normalised internal variance for non-sensing cells is now given by,

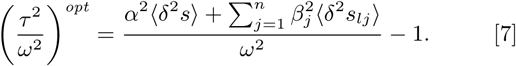

This result is similar to the results of Bull (1) and Haccou & Iwasa (2). The total variance in the environment, defined in those papers as a single parameter, equals 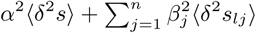 in our extended formalism. Cells should introduce noise when the relevant variance in the environment exceeds the width of the cellular fitness function.

In the appendix we evaluate the fitness difference between optimally sensing and optimally non-sensing cells, i.e., 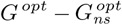. There we show (in generalised form) that 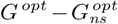 is minimally 0, which only occurs when *α* 〈*δ*^2^*s*) = 0 or *ω* → ∞. These limits are in agreement with biological intuition, since the only reason not to sense a signal, when sensing does not come at any cost, is when it contains no information about the optimum or it has no fitness consequence.

### Mutual and latent information

In order to interpret the geometric mean fitness of sensing cells from an information theoretic perspective we express it in terms of mutual information (MI). First we define the MI between the signal *s* that is being sensed and the response of the cell *p*. This way of using MI is very common in the evaluation of the accuracy of signal transduction. We find for the MI, in bits, between the sensed signal *s* and the internal response *p* (see appendix) the following relation

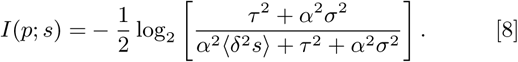

The argument of the logarithm is the fraction of all variance in *p* that is caused by internal variance (i.e., noise). The MI depends on this fraction as shown in figure 2B, which indicates that when it decreases below 10%, the relevant range of MI is approximately 1-5 bits. The relevant range of MI as a function of the internal noise depends on both the type of signal transduction and the choice of the input distribution. For instance, when two modes of a bimodal input distribution are perfectly separated and equally likely, the MI increases with 1 bit, effectively doubling the number of perceived states.

We will now introduce the crucial information measure for the fitness of sensing cells. In contrast to what is commonly done, we define an MI term between the optimum and the latent signals. From the cell’s perspective, this measure can be interpreted as ‘latent information’,

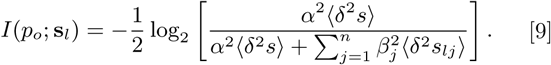

The argument of the logarithm captures the fraction of information on the optimum that the cell retrieved (figure 2C).

The relevant range of the latent information can be very low. For instance, when the sensed signal *s* contains 25% of the total information about the optimum, the latent information is 1 bit. When *s* contains 75% of all information on the optimum, the latent information is only 0.2 bit (figure 2C). An informative reference point to keep in mind is that at 0.5 bit latent information, *s* provides 50% of all information on the optimum *p*_*o*_.

Equation 9 indicates that the cell can reduce latent information by sensing more signals. By sensing an extra signal its variance is added to the numerator in equation 9, decreasing the latent information. We note that latent information is not reduced by sensing an already sensed signal more accurately.

### Maximal mutual information is not always optimal

We can write the geometric mean fitness of sensing cells (equation 4) in terms of the two mutual information measures. Before doing so, we define 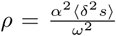, the normalised variance in optima caused by the perceived signal *s*. The geometric mean fitness of a sensing population of isogenic cells equals

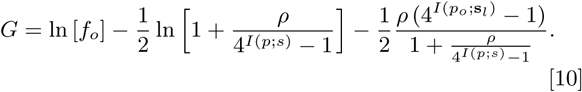

When we study *G* as a function of the mutual and latent information, given a certain value of ρ, we observe that the fitness decreases sharply in the direction of increasing latent information (figure 2D, note the low range on the x-axis).

Differentiating equation 10 with respect to the MI between *p* and *s* and setting it to 0 gives the optimal MI:

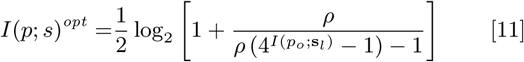

The optimal MI, *I*(*p*; *s*)^*opt*^, given a certain amount of latent information, *I*(*p*_*o*_; s_*l*_), is plotted in red in figure 2D. The fact that there is an optimum for the MI means that cells should not maximise this term. This is exactly what a great part of previous research has focused on. The optimum is valid as long as 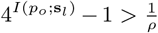, which shows that as *ρ* increases there are lower values of *I*(*p*_*o*_; s_*l*_) for which an optimal number of bits MI between *p* and *s* exists. When this optimum does not exist, fitness is maximised as *I*(*p*; *s*) approaches infinity, the fitness in this limit is given by:

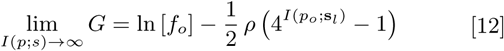

This is the maximal fitness in the ‘pure sensing’ regime, where more MI always leads to higher fitness.

### Distributed information leads to ‘noisy sensing’ of individual signals

An important conclusion can be drawn from figure 2D. It shows that for each given amount of latent information, there exists an optimal value of mutual information. This optimum converges to infinity as the latent information becomes very small.

As discussed above, the only way for cells to reduce the latent information is to sense more signals. The red line in figure 2D shows that the mutual information should increase when the latent information decreases. This concerns not just the mutual information of *p* and *s*, but also of the newly sensed signal (see appendix). This suggests that the total mutual information should increase as the cell reduces its latent information, but we do not know the optimal mutual information per signal. Within our framework, the only reason to maximise the mutual information with one signal is that this signal contains almost all information about the optimal response of the cell.

### High mutual information between one signal and a response seems rarely necessary

By distinguishing between the regimes where cells should maximise mutual information and where there exists an optimum, we can plot which strategy should be chosen for certain combinations of *ρ* and the latent information (figure 3A).

**Fig. 3.**
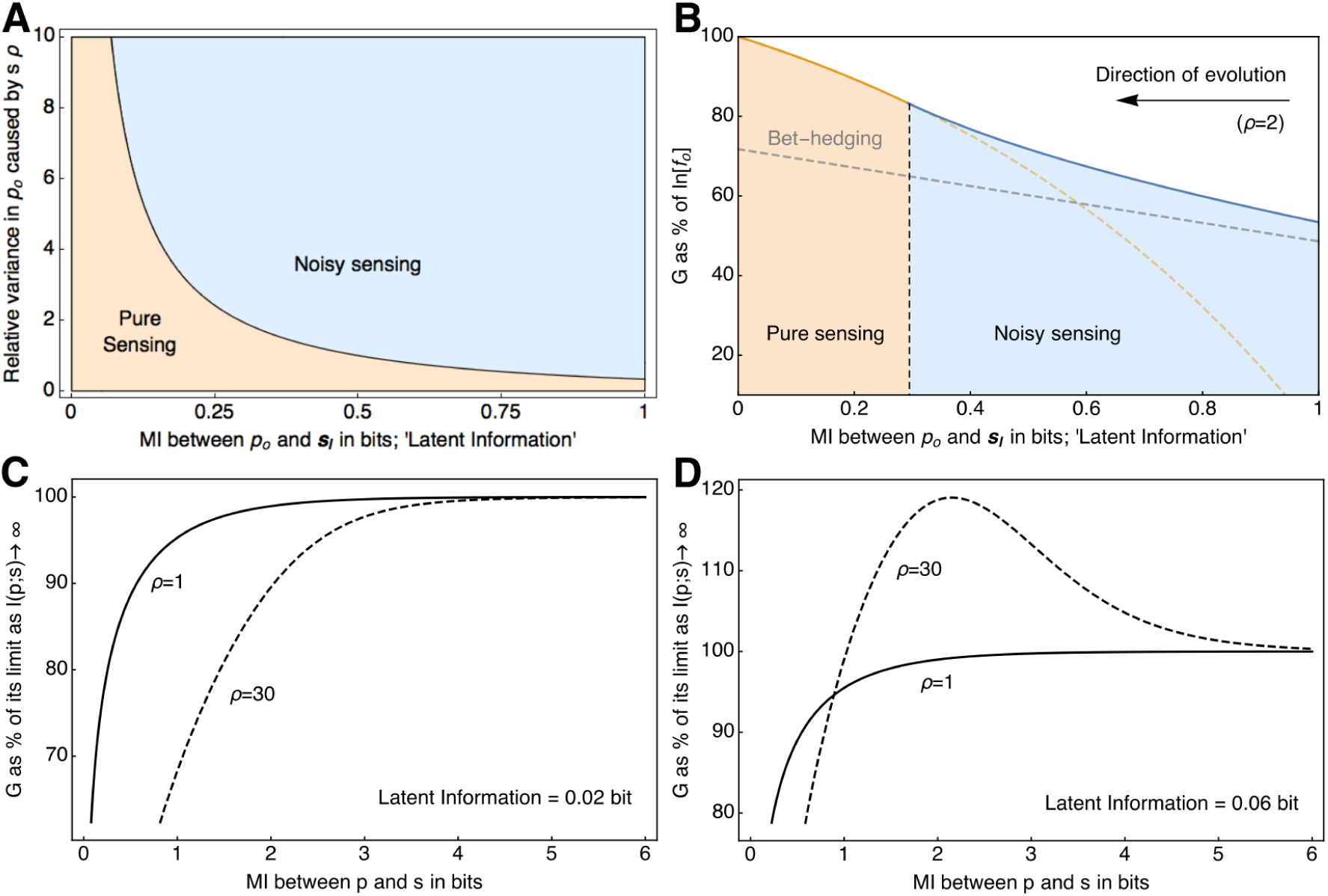
Comparison of pure and noisy sensing strategies. **A.** Optimal strategies given a certain relative variance in the optimum *p*_*o*_ caused by *s* (*ρ*) and a certain number of bits of mutual information (MI) between *p*_*o*_ and all latent signals **s**_*l*_ (i.e., the latent information). In this plot the MI between *p* and *s* is optimal in each point, so ∞ in the pure sensing regime and 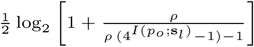 in the noisy sensing regime. **B.** The geometric mean fitness *G* as a percentage of ln [*f*_*o*_] (the maximal obtainable fitness) as a function of the MI between *p*_*o*_ and **s**_*l*_. Again the MI between *p* and *s* is optimal in each point. The dashed gray line shows the fitness of the optimal bet-hedging strategy in this situation. The direction of evolution is towards higher fitness, note that this can only be achieved by sensing more signals, as the total amount of information that is contained in one signal about the optimum is a given. **C.** *G* as a percentage of lim_*I*(*p*;*s*)→∞_ *G*, as a function of the MI between *p* an *s*. The curve saturates quickly, even when *ρ* is increased drastically. The latent information (mutual information of *p*_*o*_ and **s**_*l*_) is 0.02 bit. **D.** *G* as a percentage of lim_*I*(*p*;*s*)→∞_ *G*, as a function of the MI between *p* an *s*. As the latent information increases from 0.02 (in C) to 0.06 bit an optimum appears in the line where *ρ* = 30, meaning that in this situation cells should not maximise the MI between the sensed signal *s* and their response *p*.

Pure sensing is generally only preferred given low latent information. Another option seen in 3A, is when *ρ* becomes small. This is can be caused by two phenomena: i. *α*^2^〈*δ*^2^*s*〉 → 0, in this scenario the latent information will automatically approach infinity however (see equation 9), and ii. *ω*^2^ → ∞, now there is no consequence to missing out on the effects of the latent signals, pure sensing is now optimal because we do not consider enzyme costs to sensing, and the entire relevant fitness landscape is convex (which means averaging over a region can more often lead to a lower result than choosing one point on the landscape). How low the latent information should become in order for pure sensing to become favourable depends on the value of *ρ*.

Evolution will drive the cell to reduced latent information values (figure 3B). However, cells can only achieve this by sensing more signals. In our model the cell would therefore start sensing all relevant signals, ending up in the pure sensing regime. In addition, as long as the variance in optima caused by the sensed signal (*ρ* > 0) remains, sensing is always preferred over bet-hedging. In reality however, cells will only sense signals that contain a sufficiently large and frequent enough fitness benefit, likely creating a lower bound on the latent information. As discussed, sensing machinery with low fitness benefit might be lost via selection and genetic drift; in figure 3B we see that this could happen at high latent information.

When one signal contains almost all information about the optimum, the mutual information should be maximised (figure 3C). We note, however, that even for high *ρ* the fitness curve saturates quickly, i.e., the amount of fitness gained per bit of mutual information quickly becomes very low. This is a consequence of two factors: firstly, mutual information is an exponential measure, where the number of bits MI leads to 2^*MI*^ perceived states; secondly, the usage of normal distributions leads to 4^*MI*^ instead of 2^*MI*^, as can be seen in equation 10. In figure 3D we see how, when the latent information increases, the previously discussed optimum appears in the fitness curve. The exact latent information value when this occurs depends on the value of *ρ*.

## Discussion

Our theory indicates that a signal should be transduced as accurately as possible only when it contains nearly all information about the optimal response. When information about the optimal response is partially contained in signals that are not sensed, it is evolutionarily optimal to perceive the sensed signals noisily. When the ability to sense one signal is lost, the remaining sensed signals should not be perceived more, but *less* accurately. This is because, as the total available information about the optimal behaviour decreases, the cell should trust the available information less, and gamble more.

Maximisation of the mutual information between the input and the output of a signalling circuit has received a lot of theoretical attention. One of its advantages is that it can be applied without the use of a mechanistic, stochastic model of the signalling network. In one application, only three functions are needed: the input/output relation, the dependency of the output noise on the input, and the probability distributions of input values, i.e., of environmental states (20). Maximisation of mutual information then allows for the prediction of one of those three functions from the two others. It is often rationalised by saying that it is a requirement for fitness maximisation. Our theory has shed doubt on this argument.

How precise the signalling machinery of a single microbe should track environmental signals depends on how important those signals are for its fitness. If latent signals exist that would improve fitness if they would be sensed, then the accuracy with which sensed signals are transduced should be low. Thus, our theory predicts that the optimal mutual information between the sensed signal and the cell’s response should generally not be maximal for fitness maximisation when latent signals occur.

It is likely that latent signals occur. Firstly, because microorganisms display only a handful of signalling systems – in particular in the light of the huge number of nutrients that they can grow on. Secondly, expressing idle signalling systems reduces immediate fitness, because of biosynthetic resource consumption by non-growth promoting processes. Thirdly, idle, scanning signalling systems are evolutionary unstable; selection and drift would randomly mutate those unused systems. Natural selection for maximal geometric fitness in the presence of latent signals leads to optimal mutual information values that are not maximal.

Our approach does however have limitations, most of which are caused by our aim to create an analytically tractable model that has general applicability. We have used non-truncated normal distributions, such that distributions of compound concentrations go to minus infinity. We assumed all variances to be independent of fluctuations in compound concentrations. Also, using normal distributions might by overly simplistic. Lastly, assumptions of linearity might not always be in agreement with experimental data. However, by keeping our model analytically tractable, we obtained general qualitative insights into the fitness effects of phenotypic adaptation mechanisms by considering a minimal model. When particular systems are of interest, models that incorporate mechanisms in more detail could provide additional, system-specific insights.

It has become clear in this work that the concept of mutual information should be used with care when it comes to quantifying cell performance. Firstly, biological performance should not be measured in terms of bits, but in terms of the fitness consequences of these bits. Secondly, it is difficult to distinguish characteristics of the input distribution from characteristics of the transduction system. This aspect of mutual information can lead to a certain number of bits being perceived as low, but for the distribution of inputs that is being considered it might well be that it is close to all information that is contained in the input. This could be resolved by normalizing with the entropy of the input distribution. Thirdly, we show that the commonly used mutual information measure, the mutual information between a signal and a cell’s response, is not the relevant measure when considering its fitness consequences. In conclusion, this does not mean that mutual information cannot be used in cell biology; it only means that one has to look at what is relevant for fitness. We have shown that in cellular adaptation there is a very relevant mutual information term, which is the mutual information between all latent signals and the optimum, the latent information. Thus, it is not about how well the cell can perceive one aspect of the environment, but about how the whole environment determines the optimal behaviour. Using this perspective, it became clear that only signals that contain nearly all information about the optimal behaviour should be sensed as accurately as possible.

Our model implies that cells should improve their fitness by sensing as many informative signals as are available. In reality, we see that cells only sense a few signals. Apart from the possibility of flux-based regulation (e.g., catabolite repression (29, 30)), this is likely caused by the fact that to obtain specific information on individual signals, a cell needs separate signalling machinery for each of these signals. Whether or not a cell evolves sensing machinery for a particular signal depends on the fitness gain that can be achieved by sensing this individual signal. When this fitness gain is higher than the fitness cost of having the sensing machinery, the machinery should evolve. Even when the fitness cost of having sensing machinery is very low, sensing machinery for signals with low or infrequent fitness gain will be lost due to genetic drift. So the signals that are being sensed by microbes must have exceeded the ‘fitness threshold’, having sensing machinery for these signals is evolutionarily beneficial. The accuracy with which these signals should be sensed depends on the cumulative information in all signals that did not reach the fitness threshold, and are thus not being sensed, i.e., the latent information. This leads to the somewhat counter-intuitive conclusion that when the ability to sense one signal is lost, the other sensed signals should be perceived less accurately – not more.

This work contributes to a better understanding of optimal phenotypic adaptation strategies and of the use of information theoretic concepts. By considering multiple signals in the light of their fitness consequence we were able to show that not the mutual information between one signal and one response is what is crucial to cells, but that the latent information is what ultimately determines evolutionary success.

## ACKNOWLEDGMENTS

We would like to thank Riccardo Muolo and Daan de Groot for their time and valuable comments during our weekly discussions, and Bram van de Putte, Tom Clement and Joanne Preuter for their support and insights over many cups of coffee.

