## Appendix for "Poor sensing maximises microbial fitness when few out of many signals are sensed"

**This PDF file includes:**

Supplementary text
SI References

### Supporting Information Text

### Model derivations

For ease of comparison this appendix is structured in the same order as the results section.

**The geometric fitness model.** To determine the fitness of a sensing population we start by considering the fitness of a single cell. We assume that a maximal fitness exists for each cell, expressed in terms of the optimal concentration of one protein. The behaviour of the cell is determined by the actual concentration of this protein, $p$.

The optimal concentration of $p$ is denoted $p_o$, and is assumed to be a deterministic function $q(\mathbf{s}, \mathbf{s}_l)$. This function depends on all sensed signals, whose bulk concentrations are the components of $\mathbf{s}$, and on all so-called 'latent signals' whose concentrations are the elements of $\mathbf{s}_l$. A latent signal can be any measurable physical quantity, e.g., a nutrient concentration, in the cell's environment that the cell is not sensing, but that could be sensed by it, were the cell to change its genotype and evolve a sensing circuit. The influence of $\mathbf{s}_l$ on the optimal protein concentration is therefore defined as the concentration of $p$ the cell should have had, if it would have had the required signalling circuits for estimation of the concentrations of these latent signals. For the sake of analytical tractability we assume the optimal concentration $p_o$ to be a weighted linear sum of all signals:

$$p_o = q(\mathbf{s}, \mathbf{s}_l) = \sum_{i=1}^{k} \alpha_i s_i + \sum_{j=1}^{N-k} \beta_j s_{lj}. \quad [1]$$

Here $N$ is the total number of signals and $k$ is the number of sensed signals. The sensing cell bases its concentration $p$ on one or more externally sensed signals. Since the cell infers $s_i$ by sensing single molecules it will not know $s_i$ exactly. It will make an estimate, $s_i'$, which is assumed to follow

$$s_i' = s_i + \xi_i, \quad [2]$$

where $\xi_i$ is the Gaussian noise added due to sensing of signal $s_i$ using biochemical machinery (1, 2), i.e., a random variable with a mean of 0 and variance $\sigma_i^2$. For every signal this leads to a normally distributed $s_i'$ under fixed $s_i$: $s_i'|s_i \sim \mathcal{N}\left(s_i, \sigma_i^2\right)$. We assume all signals to be independent and can therefore also write: $\mathbf{s}'|\mathbf{s} \sim \mathcal{N}(\mathbf{s}, \Sigma)$, where $\Sigma$ is a diagonal matrix with entries $\sigma_1^2, \ldots, \sigma_k^2$. Similarly, we assume that in each single cell $p$ is based on $s_i'$ with the addition of Gaussian noise $\zeta$ due to signal transduction:

$$p = \sum_{i=1}^{k} \alpha_i s_i' + c + \zeta. \quad [3]$$

Note that we assume that $p$ follows the same linear relation to the sensed signals as the optimum does (equation 1), as we wish cells to be able to behave optimally at least in principle. The constant $c$ provides the freedom of a basal expression of $p$. The intrinsic noise $\zeta$ in the concentration of $p$ has a variance $\tau^2$. In summary, we obtain $p|\mathbf{s}' \sim \mathcal{N}\left(\sum_{i=1}^{k} \alpha_i s_i' + c, \tau^2\right)$.

We define the fitness of one cell as follows,

$$f(\mathbf{s}, \mathbf{s}_l, p) = f_o(\mathbf{s}, \mathbf{s}_l) \exp\left\{-\frac{1}{2}\frac{(p - q(\mathbf{s}, \mathbf{s}_l))^2}{\omega^2}\right\}. \quad [4]$$

The function $f_o(\mathbf{s}, \mathbf{s}_l)$ determines the maximal fitness, when $p = p_o = q(\mathbf{s}, \mathbf{s}_l)$. It is a deterministic function of $\mathbf{s}$ and $\mathbf{s}_l$, which we do not specify, and the choice of which does not change the outcomes of the analyses. The parameter $\omega$ sets the width of the fitness function: the higher its value the more forgiving the environment is (meaning optimality deviations have smaller fitness consequences).

To determine the average growth of an isogenic population in a fixed environment we need to average over its phenotypes, which have different concentrations $p$. We first determine the expected distribution of $p$ over the whole population using the definitions of the distributions for $\mathbf{s}'|\mathbf{s}$ and $p|\mathbf{s}'$ from above. The probability density functions are denoted $h_{\mathbf{S}'|\mathbf{S}}(\mathbf{s}'|\mathbf{s})$ and $g_{P|\mathbf{S}'}(p|\mathbf{s}')$ respectively. For the probability density function $g_{P|\mathbf{S}}(p|\mathbf{s})$ we obtain that $p|\mathbf{s} \sim \mathcal{N}\left(\sum_{i=1}^{k} \alpha_i s_i + c, \tau^2 + \sum_{i=1}^{k} \alpha_i^2 \sigma_i^2\right)$, i.e.

$$g_{P|\mathbf{S}}(p|\mathbf{s}) = \int g_{P|\mathbf{S}'}(p|\mathbf{s}') h_{\mathbf{S}'|\mathbf{S}}(\mathbf{s}'|\mathbf{s}) d\mathbf{s}' = \frac{1}{\sqrt{2\pi\left(\tau^2 + \sum_{i=1}^{k} \alpha_i^2 \sigma_i^2\right)}} \exp\left\{-\frac{1}{2}\frac{\left(p - \sum_{i=1}^{k} \alpha_i s_i - c\right)^2}{\tau^2 + \sum_{i=1}^{k} \alpha_i^2 \sigma_i^2}\right\}. \quad [5]$$

We can calculate the average fitness of the population over all concentrations $p$ (all phenotypes) under fixed $\mathbf{s}$ and $\mathbf{s}_l$ by multiplying $f(\mathbf{s}, \mathbf{s}_l, p)$ (eq. 4) with $g_{P|\mathbf{S}}(p|\mathbf{s})$ and integrating over $p$:

$$\begin{aligned} F(\mathbf{s}, \mathbf{s}_l) =& \langle f(\mathbf{s}, \mathbf{s}_l, p)\rangle_p = \int f(\mathbf{s}, \mathbf{s}_l, p) g_{P|\mathbf{S}}(p|\mathbf{s}) dp \\ =& \frac{f_o(\mathbf{s}, \mathbf{s}_l)\, \omega}{\sqrt{\omega^2 + \tau^2 + \sum_{i=1}^{k} \alpha_i^2 \sigma_i^2}} \exp\left\{-\frac{1}{2}\frac{\left(\sum_{i=1}^{k} \alpha_i s_i + c - q(\mathbf{s}, \mathbf{s}_l)\right)^2}{\omega^2 + \tau^2 + \sum_{i=1}^{k} \alpha_i^2 \sigma_i^2}\right\}. \end{aligned} \quad [6]$$

We have now obtained the mean fitness of the genotype in that environment.

Next, we average the fitness over a sequence of environments. As the population fitness at any point is a product of the fitnesses in previous environments (and not the sum) we should consider a geometric mean. In our framework it is most convenient to consider the sum of the logarithms of these terms. We take the natural logarithm of equation 6 and then calculate its expected value over $\mathbf{s}$ and $\mathbf{s}_l$. The equation we obtain gives the geometric mean of the population's fitness over all environments;

$$\begin{aligned} G =& \langle \ln\left[F(\mathbf{s},\mathbf{s}_l)\right]\rangle_{\mathbf{s},\mathbf{s}_l} = \int\int \ln\left[F(\mathbf{s},\mathbf{s}_l)\right] k_{\mathbf{S}}(\mathbf{s})d\mathbf{s}\, k_{\mathbf{S}_l}(\mathbf{s}_l)d\mathbf{s}_l \\ =& \langle \ln\left[f_o(\mathbf{s},\mathbf{s}_l)\right]\rangle_{\mathbf{s},\mathbf{s}_l} - \frac{1}{2}\ln\left[\frac{\omega^2+\tau^2+\sum_{i=1}^k \alpha_i^2\sigma_i^2}{\omega^2}\right] - \frac{1}{2}\frac{\int\int\left(\sum_{i=1}^k \alpha_i s_i + c - q(\mathbf{s},\mathbf{s}_l)\right)^2 k_{\mathbf{S}}(\mathbf{s})d\mathbf{s}\, k_{\mathbf{S}_l}(\mathbf{s}_l)d\mathbf{s}_l}{\omega^2+\tau^2+\sum_{i=1}^k \alpha_i^2\sigma_i^2} \end{aligned} \quad [7]$$

We denote the first term in equation 7 as $\langle \ln[f_o]\rangle$, the mean maximal fitness over all environments. To simplify the third term of equation 7 we use 1 and the assumption that all signals are independent and are normally distributed. Substituting $q(\mathbf{s},\mathbf{s}_l)$ gives for the numerator of the third term:

$$\int\left(c - \sum_{j=1}^{N-k}\beta_j s_{lj}\right)^2 k_{\mathbf{S}_l}(\mathbf{s}_l)d\mathbf{s}_l \quad [8]$$

The term is now independent of $\mathbf{s}$, and considering all $(N-k)$ latent signals are independent we can write them as separate integrals:

$$\int\cdots\int\left(c - \sum_{j=1}^{N-k}\beta_j s_{lj}\right)^2 \prod_{j=1}^{N-k} k_{S_{lj}}(s_{lj})ds_{lj} \quad [9]$$

Let us consider this integral for the first two latent signals:

$$\begin{aligned} &\int\int\left(c-\sum_{j=1}^{2}\beta_j s_{lj}\right)^2\prod_{j=1}^{2}k_{S_{lj}}(s_{lj})ds_{lj} \\ =&\int\int(c^2+\beta_1^2 s_{l1}^2+\beta_2^2 s_{l2}^2-2c\beta_1 s_{l1}-2c\beta_2 s_{l2}-2\beta_1 s_{l1}\beta_2 s_{l2})\prod_{j=1}^{2}k_{S_{lj}}(s_{lj})ds_{lj} \\ =&c^2+\beta_1^2\langle s_{l1}^2\rangle_{s_{l1}}+\beta_2^2\langle s_{l2}^2\rangle_{s_{l2}}-2c\beta_1\langle s_{l1}\rangle_{s_{l1}}-2c\beta_2\langle s_{l2}\rangle_{s_{l2}}-2\beta_1\beta_2\langle\langle s_{l1}s_{l2}\rangle_{s_{l1}}\rangle_{s_{l2}} \\ =&c^2+\beta_1^2\langle\delta^2 s_{l1}\rangle+\beta_1^2\langle s_{l1}\rangle^2+\beta_2^2\langle\delta^2 s_{l2}\rangle+\beta_2\langle s_{l2}\rangle^2-2c\beta_1\langle s_{l1}\rangle-2c\beta_2\langle s_{l2}\rangle-2\beta_1\beta_2\langle s_{l1}\rangle\langle s_{l2}\rangle \\ =&(c-\beta_1\langle s_{l1}\rangle-\beta_2\langle s_{l2}\rangle)^2+\beta_1^2\langle\delta^2 s_{l1}\rangle+\beta_2^2\langle\delta^2 s_{l2}\rangle \end{aligned} \quad [10]$$

This term has a minimum at $c=\beta_1\langle s_{l1}\rangle+\beta_2\langle s_{l2}\rangle$, corresponding to maximizing 7. Under this assumption, 10 reduces to

$$\beta_1^2\langle\delta^2 s_{l1}\rangle+\beta_2^2\langle\delta^2 s_{l2}\rangle, \quad [11]$$

which, for more than two signals, generalizes to:

$$\sum_{j=1}^{N-k}\beta_j^2\langle\delta^2 s_{lj}\rangle. \quad [12]$$

We obtain for the geometric mean fitness of an isogenic population sensing $k$ out of a total of $N$ signals:

$$G=\langle\ln[f_o]\rangle-\frac{1}{2}\ln\left[1+\frac{\tau^2+\sum_{i=1}^k\alpha_i^2\sigma_i^2}{\omega^2}\right]-\frac{1}{2}\frac{\sum_{j=1}^{N-k}\beta_j^2\langle\delta^2 s_{lj}\rangle}{\omega^2+\tau^2+\sum_{i=1}^k\alpha_i^2\sigma_i^2}. \quad [13]$$

This is a generalized equivalent of equation 3 in the main text.

To obtain the optimal internal noise equate the derivative of 13 with respect to $\tau^2+\sum_{i=1}^k\alpha_i^2\sigma_i^2$ to zero, to find the optimal relative internal variance,

$$\left(\frac{\tau^2+\sum_{i=1}^k\alpha_i^2\sigma_i^2}{\omega^2}\right)^{opt}=\begin{cases}\frac{\sum_{j=1}^{N-k}\beta_j^2\langle\delta^2 s_{lj}\rangle}{\omega^2}-1 & \text{if } \sum_{j=1}^{N}\beta_j^2\langle\delta^2 s_{lj}\rangle>\omega^2,\\ 0 & \text{if } \sum_{j=1}^{N}\beta_j^2\langle\delta^2 s_{lj}\rangle\leq\omega^2,\end{cases} \quad [14]$$

This equation tells us that when the variance in the optimal protein level, due to not knowing the values of the latent signals, exceeds the width parameter of the fitness function, it is optimal for cells to display noisy signalling.

**Sensing vs non-sensing cells.** It is clear how equation 13 changes in the two limiting cases, when $k = 0$ or when $k = N$:

$$G(k=0) = \langle \ln [f_o] \rangle - \frac{1}{2} \ln \left[ 1 + \frac{\tau^2}{\omega^2} \right] - \frac{1}{2} \frac{\sum_{j=1}^{N} \beta_j^2 \langle \delta^2 s_{lj} \rangle}{\omega^2 + \tau^2}, \qquad [15]$$

$$G(k=N) = \langle \ln [f_o] \rangle - \frac{1}{2} \ln \left[ 1 + \frac{\tau^2 + \sum_{i=1}^{N} \alpha_i^2 \sigma_i^2}{\omega^2} \right]. \qquad [16]$$

The first represents a non-sensing population (corresponding to equation **??** in the main text) and the second a population which senses all available signals, i.e., there are no latent signals anymore. The optimal internal noise levels in these cases are, respectively:

$$\left( \frac{\tau^2}{\omega^2} \right)^{opt} = \begin{cases} \frac{\sum_{j=1}^{N} \beta_j^2 \langle \delta^2 s_{lj} \rangle}{\omega^2} - 1 & \text{if } \sum_{j=1}^{N} \beta_j^2 \langle \delta^2 s_{lj} \rangle > \omega^2, \\ 0 & \text{if } \sum_{j=1}^{N} \beta_j^2 \langle \delta^2 s_{lj} \rangle \leq \omega^2, \end{cases} \qquad [17]$$

$$\left( \frac{\tau^2 + \sum_{i=1}^{N} \alpha_i^2 \sigma_i^2}{\omega^2} \right)^{opt} = 0. \qquad [18]$$

In the latter case the optimal relative internal noise is 0 because $G$ is a strictly decreasing function for all positive noise levels, so $G$ is maximal at noise level equal to 0. This is what we call a 'pure sensing' regime. Even for the non-sensing population 0 internal noise can be optimal, but only when $\sum_{j=1}^{N} \beta_j^2 \langle \delta^2 s_{lj} \rangle \leq \omega^2$.

From equations 15 and 16 it is clear that under optimised internal noise levels, the population which senses all available signals will have a higher geometric mean fitness than the non-sensing population. Only when $\sum_{j=1}^{N} \beta_j^2 \langle \delta^2 s_{lj} \rangle = 0$ both of these populations would obtain the maximal fitness $\langle \ln [f_o] \rangle$, but this would mean there is no information to be obtained from the environment. This result agrees with intuition as there are no protein costs to sensing within our framework. We can investigate the intermediate cases (with $k$ ranging from 1 to $N-1$) by considering all $\alpha_i$ and $\sigma_i$ to be equal for any $i$, and all $\beta_j$ and $\langle \delta^2 s_{lj} \rangle$ to be equal for any $j$. From equation 13 we then obtain:

$$G = \langle \ln [f_o] \rangle - \frac{1}{2} \ln \left[ \frac{\omega^2 + \tau^2 + k\,\alpha^2 \sigma^2}{\omega^2} \right] - \frac{1}{2} \frac{(N-k)\beta^2 \langle \delta^2 s_l \rangle}{\omega^2 + \tau^2 + k\,\alpha^2 \sigma^2}. \qquad [19]$$

Calculating $G(k+1) - G(k)$ gives:

$$G(k+1) - G(k) = \frac{1}{2} \left( \ln \left[ \frac{\omega^2 + \tau^2 + k\,\alpha^2 \sigma^2}{\omega^2 + \tau^2 + (k+1)\,\alpha^2 \sigma^2} \right] + \frac{(N-k)\beta^2 \langle \delta^2 s_l \rangle}{\omega^2 + \tau^2 + k\,\alpha^2 \sigma^2} - \frac{(N-k-1)\beta^2 \langle \delta^2 s_l \rangle}{\omega^2 + \tau^2 + (k+1)\,\alpha^2 \sigma^2} \right), \qquad [20]$$

which already shows that for $\omega^2 \to \infty$ nothing the population does will have a fitness consequence. Now let us consider that in both cases the population has optimized its internal noise, such that we can calculate $G(k+1)^{opt} - G(k)^{opt}$. For sensing an extra signal in a noisy sensing regime and in a pure sensing regime we obtain respectively:

$$G(k+1)^{opt} - G(k)^{opt} = \begin{cases} \frac{1}{2} \ln [\frac{N-k}{N-k-1}] & \text{if } (N-k-1)\beta^2 \langle \delta^2 s_l \rangle > \omega^2, \\ \frac{1}{2} \frac{\beta^2 \langle \delta^2 s_l \rangle}{\omega^2} & \text{if } (N-k)\beta^2 \langle \delta^2 s_l \rangle \leq \omega^2, \end{cases} \qquad [21]$$

which are both positive for any $k < N$ and as long as $\beta^2 \langle \delta^2 s_l \rangle > 0$. There is one more comparison that can be made for sensing an extra signal, which is in the transition from a noisy to a pure sensing regime, i.e. when $(N-k)\beta^2 \langle \delta^2 s_l \rangle > \omega^2 \geq (N-k-1)\beta^2 \langle \delta^2 s_l \rangle$. In this case we obtain:

$$G(k+1)^{opt} - G(k)^{opt} = \frac{1}{2} \left( \ln \left[ \frac{(N-k)\beta^2 \langle \delta^2 s_l \rangle}{\omega^2} \right] + 1 - \frac{(N-k-1)\beta^2 \langle \delta^2 s_l \rangle}{\omega^2} \right), \qquad [22]$$

which is also positive for $k < N$ and $\beta^2 \langle \delta^2 s_l \rangle > 0$.

From this we can conclude that when cells have optimised noise in signal transduction, within our model, the only reasons not to sense an extra signal are that the signal of interest contains no information on the optimum, or sensing the signal has no fitness consequence. This result is to be expected when sensing with any desired accuracy is possible without costs.

**Expressing noise in terms of mutual information.** We first define the MI between the signals that are being sensed $\mathbf{s}$ and the response of the cell $p$. Following the definition of MI (3, 4) we obtain

$$I(p; \mathbf{s}) = \mathcal{H}(p) - \langle \mathcal{H}(p \,|\, \mathbf{s}) \rangle_{\mathbf{s}}, \qquad [23]$$

where $\mathcal{H}$ denotes the entropy. Note this is the joint MI in a multiple-access channel (4). We know from equation 5 that $p|\mathbf{s} \sim \mathcal{N}\left( \sum_{i=1}^{k} \alpha_i s_i + c, \tau^2 + \sum_{i=1}^{k} \alpha_i^2 \sigma_i^2 \right)$ and given the multivariate normal distribution of the sensed signals $\mathbf{s} \sim \mathcal{N}(\langle \mathbf{s} \rangle, \Sigma)$, where $\langle \mathbf{s} \rangle$ is a vector whose components are the signal averages $\langle s_1 \rangle, \ldots, \langle s_k \rangle$ and $\Sigma$ is a diagonal matrix containing signal variances $\langle \delta^2 s_1 \rangle, \ldots, \langle \delta^2 s_k \rangle$, we can obtain the probability density function of p:

$$g_P(p) = \int g_{P|\mathbf{S}}(p|\mathbf{s}) h_{\mathbf{S}}(\mathbf{s}) d\mathbf{s} = \frac{\exp\left\{-\frac{1}{2}\frac{\left(p - c - \sum_{i=1}^{k} \alpha_i \langle s_i \rangle\right)^2}{\tau^2 + \sum_{i=1}^{k} \alpha_i^2 \langle \delta^2 s_i \rangle + \alpha_i^2 \sigma_i^2}\right\}}{\sqrt{2\pi\left(\tau^2 + \sum_{i=1}^{k} \alpha_i^2 \langle \delta^2 s_i \rangle + \alpha_i^2 \sigma_i^2\right)}}. \qquad [24]$$

Using the entropy of a normal distribution, we insert the probability density functions of $p$ and $p|\mathbf{s}$ in equation 23:

$$\begin{aligned} I(p;\mathbf{s}) =& \frac{1}{2}\log_2\left[2\pi e\left(\tau^2 + \sum_{i=1}^{k} \alpha_i^2 \langle \delta^2 s_i \rangle + \alpha_i^2 \sigma_i^2\right)\right] - \frac{1}{2}\log_2\left[2\pi e\left(\tau^2 + \sum_{i=1}^{k} \alpha_i^2 \sigma_i^2\right)\right] \\ =& -\frac{1}{2}\log_2\left[\frac{\tau^2 + \sum_{i=1}^{k} \alpha_i^2 \sigma_i^2}{\tau^2 + \sum_{i=1}^{k} \alpha_i^2 \langle \delta^2 s_i \rangle + \alpha_i^2 \sigma_i^2}\right]. \end{aligned} \qquad [25]$$

The argument of the logarithm in equation 25 is the fraction of all variance in $p$ that is caused by internal variance (i.e., noise). Note that this can also be written in the more familiar form

$$I(p;\mathbf{s}) = \frac{1}{2}\log_2\left[1 + \frac{\sum_{i=1}^{k} \alpha_i^2 \langle \delta^2 s_i \rangle}{\tau^2 + \sum_{i=1}^{k} \alpha_i^2 \sigma_i^2}\right], \qquad [26]$$

where $\frac{\sum_{i=1}^{k} \alpha_i^2 \langle \delta^2 s_i \rangle}{\tau^2 + \sum_{i=1}^{k} \alpha_i^2 \sigma_i^2}$ is the signal to noise ratio. For our purpose the notation in equation 25 will prove to be more intuitive.

**Defining latent information: mutual information between the optimum and all latent signals.** We can derive the MI between the optimum $p_o$ and all latent signals $\mathbf{s}_l$ in a manner similar to how we derived the MI between $p$ and $\mathbf{s}$ above. This measure is what we call the 'latent information'. As $p_o$ is the sum of independent normally distributed random variables, as shown in equation 1, and having assumed all signals to be independent, we obtain a normally distributed $p_o$ with the following mean and variance respectively:

$$\langle p_o \rangle = \sum_{i=1}^{k} \alpha_i \langle s_i \rangle + \sum_{j=1}^{N-k} \beta_j \langle s_{lj} \rangle, \qquad [27]$$

$$\langle \delta^2 p_o \rangle = \sum_{i=1}^{k} \alpha_i^2 \langle \delta^2 s_i \rangle + \sum_{j=1}^{N-k} \beta_j^2 \langle \delta^2 s_{lj} \rangle. \qquad [28]$$

So its probability density function is given by:

$$q_{P_o}(p_o) = \frac{\exp\left\{-\frac{1}{2}\frac{\left(p_o - \sum_{i=1}^{k} \alpha_i \langle s_i \rangle - \sum_{j=1}^{N-k} \beta_j \langle s_{lj} \rangle\right)^2}{\sum_{i=1}^{k} \alpha_i^2 \langle \delta^2 s_i \rangle + \sum_{j=1}^{N-k} \beta_j^2 \langle \delta^2 s_{lj} \rangle}\right\}}{\sqrt{2\pi\left(\sum_{i=1}^{k} \alpha_i^2 \langle \delta^2 s_i \rangle + \sum_{j=1}^{N-k} \beta_j^2 \langle \delta^2 s_{lj} \rangle\right)}}. \qquad [29]$$

It is clear that when $\mathbf{s}_l$ is fixed, the variances of the latent signals $\langle \delta^2 s_{l_j} \rangle$ will be 0:

$$q_{P_o|\mathbf{S}_l}(p_o|\mathbf{s}_l) = \frac{\exp\left\{-\frac{1}{2}\frac{\left(p_o - \sum_{i=1}^{k} \alpha_i \langle s_i \rangle - \sum_{j=1}^{N-k} \beta_j \langle s_{lj} \rangle\right)^2}{\sum_{i=1}^{k} \alpha_i^2 \langle \delta^2 s_i \rangle}\right\}}{\sqrt{2\pi \sum_{i=1}^{k} \alpha_i^2 \langle \delta^2 s_i \rangle}}. \qquad [30]$$

Now we can make use of the probability density functions defined in equations 29 and 30 and the entropy of a normal distribution, to calculate the MI between $p_o$ and $\mathbf{s}_l$:

$$\begin{aligned} I(p_o;\mathbf{s}_l) =& \mathcal{H}\left(p_o\right) - \langle \mathcal{H}\left(p_o\,|\mathbf{s}_l\,\right)\rangle_{\mathbf{s}_l} \\ =& \frac{1}{2}\log_2\left[2\pi e\left(\sum_{i=1}^{k}\alpha_i^2\langle\delta^2 s_i\rangle + \sum_{j=1}^{N-k}\beta_j^2\langle\delta^2 s_{lj}\rangle\right)\right] - \frac{1}{2}\log_2\left[2\pi e\sum_{i=1}^{k}\alpha_i^2\langle\delta^2 s_i\rangle\right] \\ =& -\frac{1}{2}\log_2\left[\frac{\sum_{i=1}^{k}\alpha_i^2\langle\delta^2 s_i\rangle}{\sum_{i=1}^{k}\alpha_i^2\langle\delta^2 s_i\rangle + \sum_{j=1}^{N-k}\beta_j^2\langle\delta^2 s_{lj}\rangle}\right]. \end{aligned} \qquad [31]$$

Here the fraction inside the logarithm shows how much information the cell has on the optimum compared to all information that determines the optimum. By sensing an extra signal this signal's variance is added in the numerator in equation 31, decreasing the latent information.

**Writing the fitness of sensing cells in terms of mutual and latent information.** Using the MI between $p$ and $\mathbf{s}$ (equation 25) and the MI between $p_o$ and $\mathbf{s}_l$ (equation 31, the latent information), we can write the geometric mean fitness (equation 13) in terms of these measures. We first rewrite equation 25 to derive:

$$\tau^2 + \sum_{i=1}^{k}\alpha_i^2\sigma_i^2 = \frac{\sum_{i=1}^{k}\alpha_i^2\langle\delta^2 s_i\rangle}{4^{I(p;\mathbf{s})}-1}. \qquad [32]$$

And similarly from equation 31,

$$\sum_{j=1}^{N-k}\beta_j^2\langle\delta^2 s_{lj}\rangle = \sum_{i=1}^{k}\alpha_i^2\langle\delta^2 s_i\rangle(4^{I(p_o;\mathbf{s}_l)}-1). \qquad [33]$$

Now we define $\rho = \frac{\sum_{i=1}^{k}\alpha_i^2\langle\delta^2 s_i\rangle}{\omega^2}$, which is a measure for the variance in the optimum caused by the perceived signals $\mathbf{s}$ (i.e., the total information readily available to the cell), relative to the fitness width. Inserting our results from equations 32 and 33 in equation 13 gives:

$$\begin{aligned} G =& \ln\left[f_o\right] - \frac{1}{2}\ln\left[1 + \frac{\sum_{i=1}^{k}\alpha_i^2\langle\delta^2 s_i\rangle}{\omega^2 4^{I(p;\mathbf{s})}-1}\right] - \frac{1}{2}\frac{\sum_{i=1}^{k}\alpha_i^2\langle\delta^2 s_i\rangle(4^{I(p_o;\mathbf{s}_l)}-1)}{\omega^2 + \frac{\sum_{i=1}^{k}\alpha_i^2\langle\delta^2 s_i\rangle}{4^{I(p;\mathbf{s})}-1}} \\ =& \ln\left[f_o\right] - \frac{1}{2}\ln\left[1 + \frac{\rho}{4^{I(p;\mathbf{s})}-1}\right] - \frac{1}{2}\frac{\rho\left(4^{I(p_o;\mathbf{s}_l)}-1\right)}{1+\frac{\rho}{4^{I(p;\mathbf{s})}-1}} \end{aligned} \qquad [34]$$

Differentiating equation 34 with respect to the MI between $p$ and $\mathbf{s}$ and setting it to 0 gives the optimal MI:

$$I(p;\mathbf{s})^{opt} = \frac{1}{2}\log_2\left[1 + \frac{\rho}{\rho\left(4^{I(p_o;\mathbf{s}_l)}-1\right)-1}\right] \qquad [35]$$

This optimum is valid as long as $4^{I(p_o;\mathbf{s}_l)} - 1 > \frac{1}{\rho}$, which shows that as $\rho$ increases there are lower values of $I(p_o;\mathbf{s}_l)$ for which an optimal number of bits MI between $p$ and $\mathbf{s}$ exists. When this optimum does not exist fitness is maximised as $I(p;\mathbf{s})$ approaches infinity, the fitness in this limit is given by:

$$\lim_{I(p;\mathbf{s})\to\infty} G = \ln\left[f_o\right] - \frac{1}{2}\rho\left(4^{I(p_o;\mathbf{s}_l)}-1\right) \qquad [36]$$

This is the maximal fitness in the 'pure sensing' regime, where more MI always leads to higher fitness.

**Exploring the mutual information through individual sensory pathways.** In equation 25 we have given the mutual information between all sensed signals and the response protein. We can wonder how this relates to the mutual information between each individual signal and the optimum. Each 'sensing channel' has a certain capacity which is given by

$$I(p;s_m) = -\frac{1}{2}\log_2\left[\frac{\tau^2+\alpha_m^2\sigma_m^2}{\tau^2+\sum_{i=1}^{k}\left(\alpha_i^2\langle\delta^2 s_i\rangle+\alpha_i^2\sigma_i^2\right)}\right]. \qquad [37]$$

We can see that if we would sum this equation over all signals we would get a sum of logarithms. This term, $\sum_{i=1}^{k} I(p;s_i)$, is always larger than the mutual information between all signals and the response protein simultaneously (i.e., the joint mutual information) (4):

$$\sum_{i=1}^{k} I(p;s_i) > I(p;\mathbf{s}). \qquad [38]$$

This is quite an intuitive result, considering the information between the set of signals and the response protein could never exceed the summed information capacity of all individual channels.

Unfortunately this does not directly provide us with a bound on the optimal mutual information between each individual signal and the response protein. We do know that in the optimum equation 14 must hold. Remember that we concluded that as long as $\sum_{j=1}^{N-k}\beta_j^2\langle\delta^2 s_{lj}\rangle > \omega^2$, there is an optimum for $\tau^2+\sum_{i=1}^{k}\alpha_i^2\sigma_i^2$ which is larger than zero. This means that in these circumstances it would be suboptimal to have infinite information capacity in all channels. Theoretically, there is no optimal distribution of the information capacity through each channel (equation 37) within our framework. However, let us consider an exponentially increasing enzyme cost of surpressing noise in each transmission channel, originating from the proportionality of noise in biochemical processes to $\frac{1}{\sqrt{n}}$, with $n$ as the number of molecules (1, 5). In this scenario it is likely that information capacity is distributed quite evenly over all channels, as this would be the most economic way of conforming to equations 14 and 35. (The precise distribution would then depend on the exact costs of the enzymes in each signalling pathway.)
